# The right time and place: time- and age-dependent vaccine-enhanced mucosal immunity to parasite infection

**DOI:** 10.1101/2021.04.28.441781

**Authors:** Wei Liu, Tom N. McNeilly, Mairi Mitchell, Stewart T.G. Burgess, Alasdair J. Nisbet, Jacqueline B. Matthews, Simon A. Babayan

## Abstract

Individuals vary broadly in their response to vaccination and subsequent exposure to infection, causing persistence of both infection and transmission. The prevalence of poor vaccine responders hampers the development of vaccines, especially against parasitic helminths. Yet despite having substantial economic and societal impact, the immune mechanisms that underlie such variability, especially at the site of parasite infection, remain poorly understood. Previous trials using a prototype vaccine for the control of the gastric parasitic *Teladorsagia circumcincta*, one of the highest impact parasites affecting sheep, revealed substantial variation in protection between individuals, which we hypothesised may in part be driven by age at vaccination. Here, to characterise how immunity at the mucosal site of infection developed in vaccinated lambs, we inserted gastric cannulae into the abomasa (true stomachs) of three-month- and six-month-old lambs before vaccination, and performed a longitudinal analysis of their local immune response during subsequent challenge infection. We found that the vaccine caused systemic changes in the baseline immune profile within the abomasum before any parasite exposure had occurred and reduced parasite burden and egg output once lambs were infected, regardless of age. However, age affected how vaccinated lambs responded to subsequent infection across multiple immune pathways, with only a minority of protective immune pathways being independent of age. This resulted in younger lambs being more susceptible to infection regardless of vaccine status. The identification of age-dependent (mostly adaptive) and age-independent (mostly innate) protective immune pathways should help refine the formulation of vaccines against these and potentially other helminth parasites of ruminants, and could indicate specificities of anti-helminth immunity more generally.

---

Individuals vary widely in their responses to vaccination and little is known about the causes underlying low vaccine efficacy, nor how effective a vaccine will be “in the real world” against the pathogen for which it has been developed. In fact, for most pathogens, imperfect vaccination is the rule rather than the exception (1). This is particularly problematic for helminths, as vaccination is the most promising alternative to anthelmintic drug treatments (2) in the face of extensive drug resistance, which is rising in human parasites (3, 4) and ubiquitous in many helminths of livestock, including in sheep (5–7). In both vaccinated and non-vaccinated animals, even a minority of infected individuals can release substantial numbers of helminth eggs into the environment, ensuring that infection persists within populations (8). While the need for better prediction of vaccine responsiveness has long been recognised, the complexity of the factors involved — from variation in genetic background (9, 10) to differences in age, sex, and immune history (11, 12), combined with immune evasion strategies deployed by helminths to ensure persistent infection (13–15) — have so far hampered the development of effective sub-unit vaccines for controlling helminths (16). Resistance to several anthelmintic compounds in sheep nematodes in the UK alone is estimated to cost in excess of £40M annually (17). With the need for effective vaccines ever more pressing, it is therefore urgent to identify which immune pathways mediate vaccine efficacy, and to understand how they change over time and with host age.

We recently developed a prototype subunit vaccine against *Teladorsagia circumcincta*, a major contributor to parasitic gastroenteritis in sheep. This parasitic nematode resides in the abomasum (the gastric compartment of the ruminant stomach) and is primarily a cause of disease in lambs. Third-stage larvae (L3) penetrate glands within 24 h of infection and grow rapidly, undergoing two moults before emerging into the abomasal lumen approximately 10 days post-infection (18). The resulting pathology manifests as anorexia, diarrhoea and poor productivity. The prototype vaccine achieves 58–70% reduction in worm burdens and up to 73–92% fewer eggs at the peak of worm egg shedding compared to challenge controls (19). Such reductions in cumulative worm faecal egg counts (cFEC) are expected to have a substantial impact on pasture contamination and modelling studies indicate that reductions of this magnitude will parallel effective anthelmintic-driven parasite control measures (20). However, substantial variation in the efficacy of this vaccine has been observed among individuals (19, 21). Such variability can be due to multiple genetic and environmental factors (15, 22, 23). While genetic factors can be manipulated by selective breeding and environmental factors such as nutritional resource can be optimised under farming conditions, any vaccine against *T. circumcincta* is likely to be most effective if administered to lambs before exposure to worms on pasture and to protect them during the late spring/summer period in which their growth rate is most susceptible to the impact of parasitic gastroenteritis. This, however, raises concerns since the immature immune system is known to respond poorly to immunisation (24–26). We therefore sought to compare the efficacy and the immune responses to our prototype vaccine in two different age groups, 3-month-old (3mo) and 6-month-old (6mo) lambs to attempt to understand age- and vaccine-dependent protective immunity at the site of infection to inform further optimisation of the vaccine.

How lambs of different ages respond immunologically to our prototype vaccine over the course of repeated exposure and during chronic infection by *T. circumcincta* is not known. We therefore analysed the immune gene expression pathways elicited by immunisation and subsequent repeated challenge infection with *T. circumcincta* in the abomasum using sequential gastric biopsies aligned to a novel machine-based learning approach, to characterise: (a) which immune pathways are associated with variation in parasite burdens and parasite egg output and when they are expressed during the course of infection; at what time during the course of the infection are those pathways associated with protection, and (c) how those pathways are affected by both vaccination and host age.

## Results

### Vaccination reduced worm burdens and egg shedding in both 3mo and 6mo lambs

We sought to identify the immune mechanisms underlying variation in the ability of the prototype vaccine to control *T. circumcincta* infection in lambs and to characterise how lamb age affects vaccine efficacy. Three- and six-month-old lambs were vaccinated using our previously-published protocol (19). Briefly, fifteen 3mo and fifteen 6mo lambs were administered an eight-protein cocktail of *T. circumcincta* recombinant antigens with Quil-A adjuvant, three times at three-weekly intervals. All vaccinated and non-vaccinated age-matched control lambs were then all exposed to repeated (“ trickle”) infections with 12 consecutive challenges with *T. circumcincta* infective larvae (L3) delivered in measured doses spanning four weeks to mimic field challenge conditions (Fig. S1).

Vaccination led to significant reductions in both worm burden assessed at *post mortem* (Fig. 1A; P = 0.002) and cumulative faecal egg counts (cFEC) (Fig. 1B; P = 0.026) in both age groups relative to control lambs. In addition, 6mo lambs, whether immunised or not, controlled worm burdens more effectively than 3mo lambs (P = 0.012). However, total egg output differed only marginally between age groups and was characteristically variable.

**Fig. 1.**
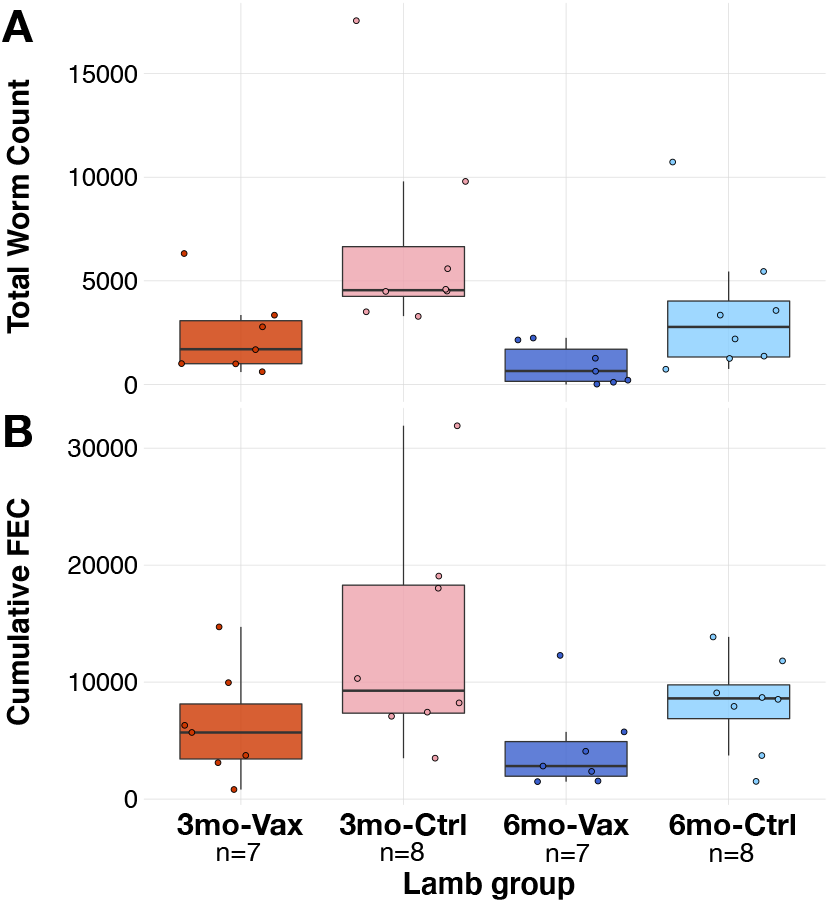
Vaccine- and age-mediated control of *T. circumcincta* infection. (*A*), Worm burdens and (*B*), total egg output in 3 month-old lambs with vaccination (3mo-Vax), 3 month-old lambs with adjuvant only (3mo-Ctrl), 6 month-old lambs with vaccination (6mo-Vax), and 6 month-old lambs with adjuvant only (6mo-Ctrl). Immunisation with the prototype vaccine led to a median 73.7% reduction (P_vacc_ = 0.002, GLM) in worm counts and 50% reduction (P_vacc_ = 0.026, GLM) in cumulative egg output.

### Pathway enrichment was most predictive of vaccine efficacy prior to challenge and following parasitological events

Because worm burdens and cFEC varied substantially between individuals regardless of age or vaccination, we then sought to identify (i) which immune pathways were associated with controlling worm burdens at post-mortem (49 days after the start of challenge infection) and cFEC, and (ii) when their expression in the abomasum best predicted these measures of parasite infection. To address both questions, we built an analysis pipeline to identify the immune pathways that best predicted infection outcomes (Fig. S2). RNAseq was performed on biopsies taken repeatedly from the abomasum at temporal intervals. These produced six transcriptomes spanning 42 days for each lamb, beginning post-vaccination but shortly before the first challenge and ending one week prior to the end of the trial when the worm burden analysis was undertaken (Fig. S1). T-distributed stochastic neighbour embedding (t-SNE) suggested there was weak structure within the gene expression read counts of all 180 whole transcriptomes (Fig. S3). We then clustered transcripts into co-expression modules regardless of age using a weighted correlation network analysis (WGCNA (27)). This generated a reduced list of variables, each of which represented a set of genes that behaved similarly across all transcriptomes. To identify which of the resulting modules contained genes of interest, we used worm burdens and cFEC as response variables in ElasticNet regression models to which we mapped the eigenvalue of each WGCNA module, and used the coefficients learned by the ElasticNet to rank the modules according to the strength and direction of their association with either worm burden or cFEC (Fig. S4).

We then extracted all genes included within the modules that predicted either worm burden or cFEC, then entered that gene list into a canonical pathway analysis, and selected the resulting immune pathways (28). Fig. 2 presents the immune pathways that robustly predicted worm burdens of cFEC and their level of enrichment (proportion of genes within specific pathways that was detected in the transcriptome) at each time-point. This revealed that most immune pathways that predicted either parasite burden 49 days (D49) post-infection or cFEC were already enriched by D0, i.e. following immunisation, but prior to the first challenge. Pathways that were significantly represented exclusively at this time-point included B cell development and activation signalling, as well as innate pathways involved in interferon and TNF signalling, macrophage stimulating protein receptor signalling (MSP-RON), and the complement system (Fig. 2, inner ring).

**Fig. 2.**
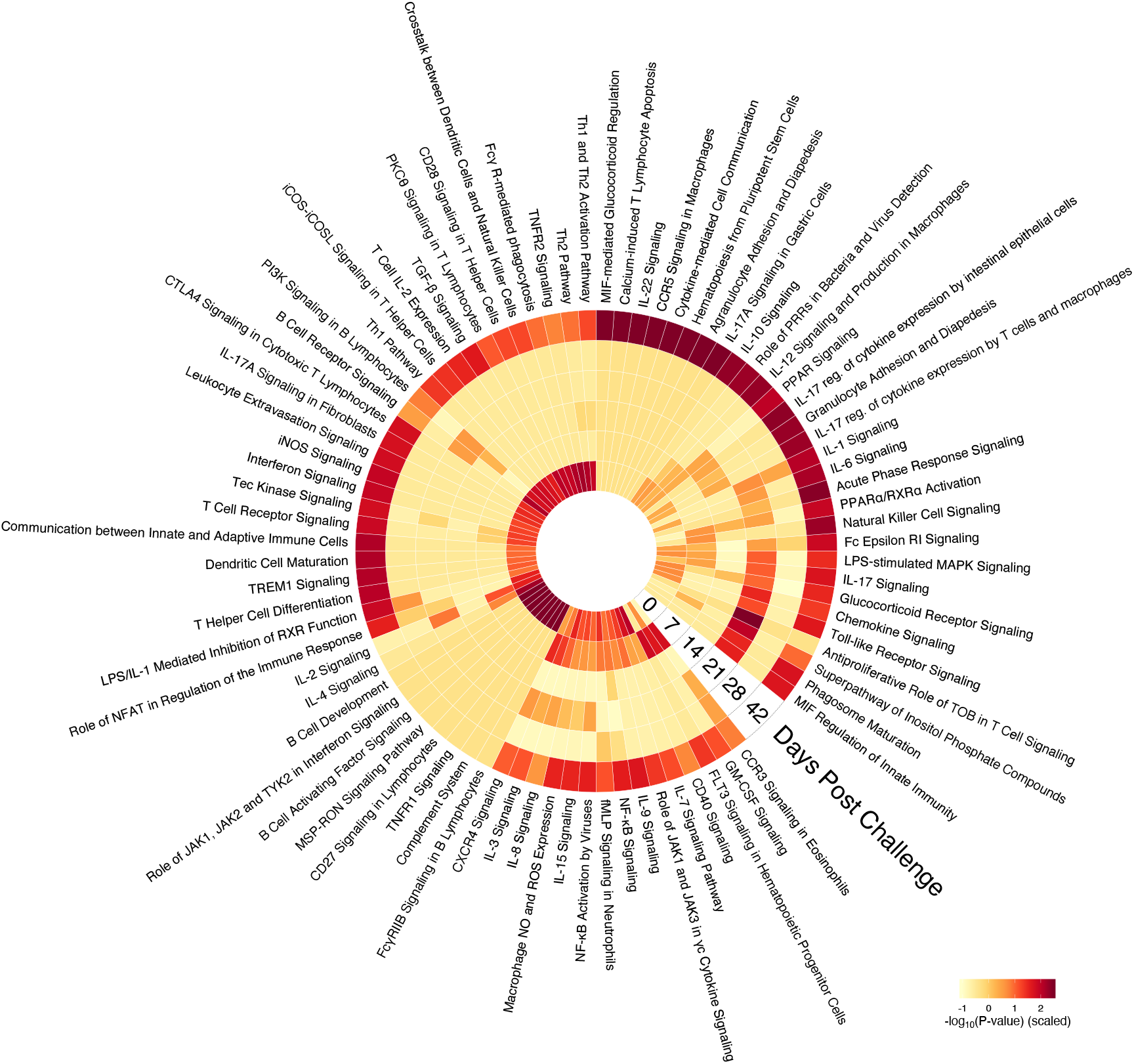
Temporal dynamics of the expression of immune pathways that predict either post-mortem worm burden or cFEC identified by supervised machine learning ElasticNet. Heat map indicates the pathway enrichment reported as negative 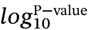, where the dark red denotes stronger enrichment within each pathway. Each concentric circle represents one of six time-points at which abomasal biopsies were taken after challenge infection. Worm burdens predicted by these pathways were measured at D49 post challenge.

All other pathways significantly enriched at D0 were also enriched at subsequent time-points. In particular, pathways whose expression correlated with worm or egg counts showed significant enrichment at days 7, 21, and 42 post-challenge, consistent with the timing of three significant phases of the life cycle, i.e. at D7, upon emergence of the first cohort of late fourth stage larvae (L4)/early adults from the gastric glands into the lumen (29, 30), coinciding with high enrichment scores of pathways involved in the regulation of innate responses (NF-κB, eosinophil CCR3, GMCSF, NO and ROS expression by macrophages, IL-2/IL-15, IL-3, and IL-8 signalling) as well as antigen-processing and lymphoid cell activation (FLT3, IL-7, CD40, CXCR4); at D21, co-incident with the initiation of egg-laying by the founding population of *T. circumcincta*, a further set of immune pathways was activated, including inflammatory and stress response signals (LPS-stimulated MAPK, MIF, glucocorticoid receptors, IL-17, and TLR signalling) and suppression of lymphocyte proliferation (anti-proliferative role of TOB in T cell signalling). Enrichment at D42 was elevated in the majority of pathways depicted in Fig. 2 (see full timecourse for all pathways in Fig. S6 and Supplementary Data 2). Being close to the post-mortem time-point (D49), this likely reflects near-contemporaneous and potentially shorter-lived correlations between immune gene expression and parasite burdens, and were thus not considered further in the prediction of protective immunity. The very low enrichment scores in the remaining time-points likely indicates either low expression of the corresponding genes due to biological regulatory processes, or too much variance for any statistical pattern to be detected. We therefore decided to further focus on time-points D0, D7, and D21 for their relevance to predicting vaccine-mediated immunity against *T. circumcincta* in these lambs.

### Direction and strength of correlation between immune pathway expression and parasitological measures varied between time-points

To identify the time-points at which the selected pathways (Fig. 2) best predicted reduced parasite burdens and cFEC, we assessed the direction and strength of the correlations between each pathway and either parasite burden (Fig. 3A) or cFEC (Fig. 3B) before (D0) or 7 and 21 days following the first exposure to *T. circumcincta* L3. For ease of interpretation, we then clustered immune pathways according to whether they were negatively associated with parasite burdens at each, or all, of these time-points.

**Fig. 3.**
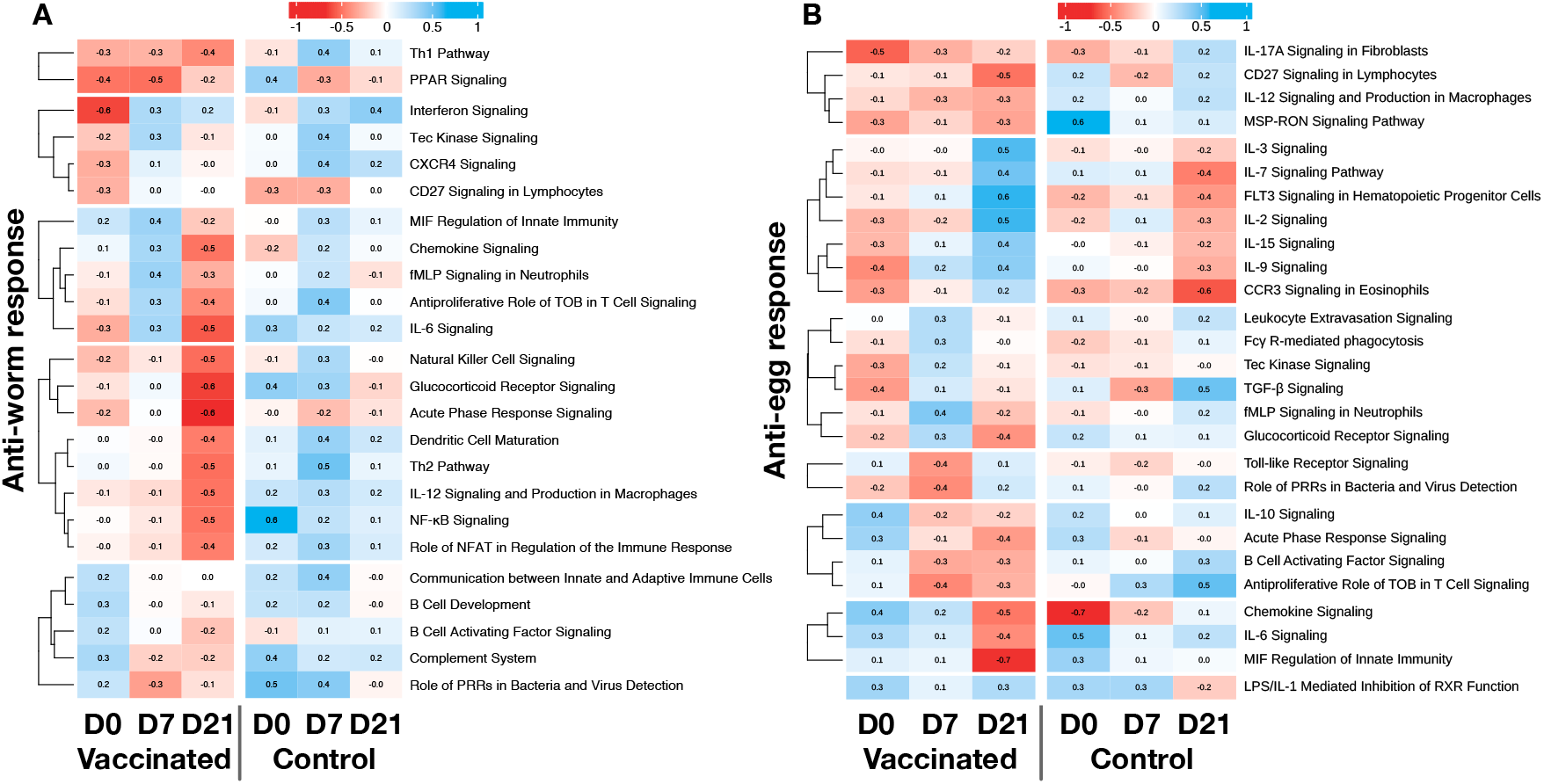
Heat map of correlation coefficients between (*A*), worm burdens or (*B*), cFEC and gene expression in the selected pathways before challenge (D0), seven (D7) and 21 (D21) days post challenge in vaccinated and control lambs. Colours represent negative (red) and positive (blue) Pearson correlation coefficients for each comparison. The immune pathways were clustered using k-means according to their correlation patterns over time in the vaccinated group.

Comparing the expression of immune pathways at D0 in vaccinated lambs and non-vaccinated controls indicated that IL-6, Th1, PPAR, and interferon signalling pathways elicited by the vaccine were instrumental in reducing parasite numbers at post-mortem (D49), while higher activation of IL-17A, IL-9, CCR3, and TGF-β signalling pathways in vaccinated lambs prior to infection predicted lower egg shedding. In non-vaccinated lambs, no pathways were predictive of lower worm numbers, whereas IL-17A, CCR3, and chemokine signalling pathways were identified as predictors of low cFEC.

At D7, Th1 and PPAR signalling remained negatively associated with post-mortem worm burden in vaccinates, while most other pathways were either positively or not associated with worm burdens. With regard to egg shedding, in vaccinated lambs activation of IL-17A remained negatively associated with cFEC, with pathways associated with regulation of T cell activation and Th1 polarisation (TOB and IL-12 pathways, respectively) also associated with reduced cFEC. Similar to D0, no immune pathways were predictive of lower parasite numbers in control lambs, although CCR3 and chemokine signalling pathways remained consistently, though weakly, negatively associated with cFEC.

Finally, at D21 vaccinated lambs differed from controls by exhibiting strong negative associations between worm burdens and chemokine signalling, IL-6, NFκB, natural killer cells, glucocorticoid receptor and acute phase response signalling, suggesting a potential role for responses to tissue injury in vaccine-induced protection against *T. circumcincta* (Fig. 3). Additionally, pathways associated with T cell activation and polarisation (IL-12, DC maturation, TOB, NFAT, Th1 and Th2 signalling pathways) were all negatively associated with worm burden at this time-point. In the non-vaccinated lambs, no pathways at D21 showed any association with worm burdens. With regard to predicting vaccine-induced impacts on cFEC, many of the pathways associated with reduced worm burdens at D21, in particular chemokine signalling, IL-6, and acute phase response signalling, were also predictive of lower cFEC in vaccinates. Lower egg shedding in vaccinated lambs was also associated with increased expression of CD27 in lymphocytes, and macrophage migration inhibitory factor (MIF). Interestingly, in non-vaccinated lambs, IL-7, IL-9, FLT3, and CCR3 signalling were negatively associated with parasite egg shedding at D21, whereas these signalling pathways were positively associated with cFEC in vaccinated lambs at the same time-point post-challenge.

### Protective immune pathways were differentially affected by age and vaccination status

We then sought to analyse how the age of lambs at immunisation affected their expression of protective immune pathways at the same time-points. Overall, the mean gene expression levels for most of the pathways described above were most strongly expressed in the older (6mo) vaccinates at D21 than at the two other time-points (Fig. 4A). Six-month-old lambs also displayed a greater differential in gene expression between D0 and D21, regardless of vaccination status. Indeed, while mean gene expression of immune pathways over all time-points was often greater in 3mo lambs, gene expression levels varied little over time in the 3mo lambs, particularity in non-vaccinated individuals. To assess how age and immunisation explained the observed variation in the expression of the focal immune pathways, we constructed generalised linear models for each pathway in turn with age, vaccine status, and the interaction between age and vaccine as explanatory variables. Most protective pathways identified by the ElasticNet (Fig.3) were affected by lamb age, especially pathways involved in the activation of adaptive immunity and its maintenance, spanning Th1, Th2, and Th17 pathways (Fig. 4B ‘Age × Vaccine’ column).

**Fig. 4.**
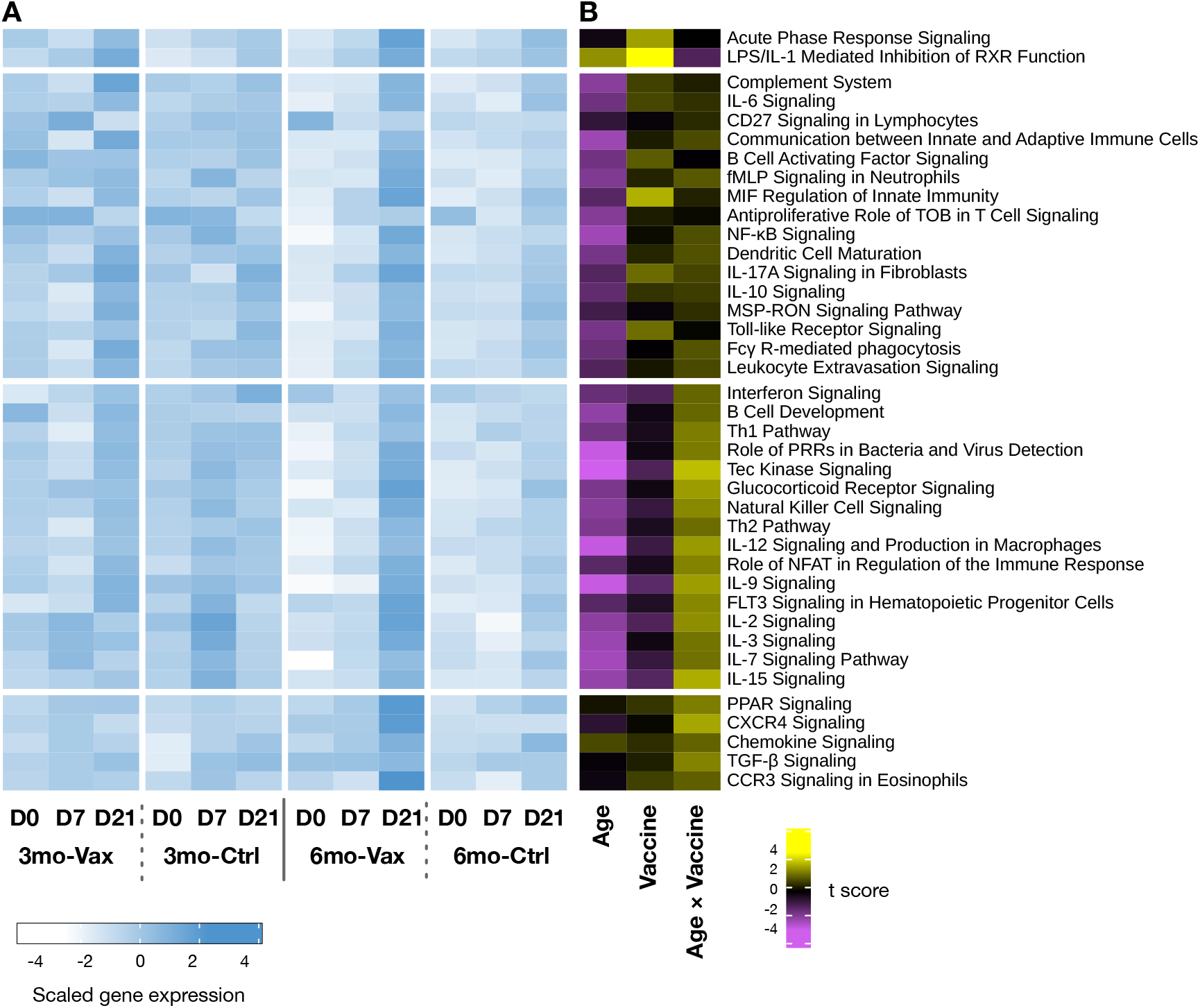
Gene expression within immune pathways significantly affected by immunisation, age, or both. (*A*) Heat map of gene expression over time within the four treatments (3mo-Vax, 3mo-Ctrl, 6mo-Vax, and 6mo-Ctrl). High to low scaled expression is denoted with blue to white hues. (*B*) Heat map of a generalised mixed model t value statistic (number of standard deviations from the mean) (31) indicating the effect of age, vaccination, and the interaction between age and vaccination on the expression of each pathway. Purple indicates a dampening effect on the expression of the pathway while yellow indicates increased expression of that pathway relative to the global average. Pathways included are those presented in Fig. 2A and 2B. The full list of gene expression is available in Fig. S6.

Pathways that were elicited by vaccination independently of age included acute phase response signalling, LPS-mediated inhibition of RXR function, MIF Regulation of Innate Immunity, and Toll-like Receptor Signalling (Fig. 4B, ‘Vaccine’ column). Finally, regardless of vaccination, only Antiproliferative Role of TOB in T Cell Signalling and possibly the Complement System were affected by age, with 6mo lambs expressing them at lower levels than 3mo lambs (Fig. 4B ‘Age’ column).

## Discussion

Responses to immunisation vary greatly between individuals. While factors such as age, sex, and nutritional status are known to modulate vaccine efficacy, substantial variation in immune responses between individuals and over time within the same individual confound efforts to develop vaccines against chronic infections. For sheep exposed to gastrointestinal parasites such as *T. circumcincta*, the immune system is a crucial line of defence as drug resistance among parasites becomes ubiquitous. Vaccines therefore have considerable potential to protect the welfare and productivity of livestock. However, which immune pathways a vaccine ought to elicit against gastrointestinal helminth infections is largely unknown and may not necessarily be the same naturally-induced responses that the hosts have evolved under normal exposure to the pathogen. Indeed, hosts may prioritise tissue integrity and tolerance over protective immunity, which may be counterproductive to livestock welfare and productivity goals. Further, mass immunisation depends on the creation of a recombinant vaccine. This has proven difficult, with few successes despite decades of research.

We have recently developed a prophylactic vaccine that shows great promise against *T. circumcincta* (19). *However, for this prototype to deliver consistent protection a better understanding of the immune mechanisms that underlie the observed imperfect immunisation in sheep of relevant ages is needed (20). The present study was aimed at characterising, among the immune pathways elicited by the vaccine before and during infection by T. circumcincta*, those that led to protection, and at which phase of the infection they were most effective. We monitored immune responses of vaccinated sheep at the site of infection (the abomasum) over the first 42 days of a trickle challenge infection, and measured parasite burdens on day 49. To assess how host age affects vaccine efficacy, we immunised lambs at 3 or 6 months of age. Consistent with our previous reports (19, 32), the vaccine led to significant reductions of both worm burdens and egg outputs compared to adjuvant (Quil A) only recipients (Fig. 1). However, 3-month-old lambs remained more heavily infected than 6-month-old lambs throughout the experiment, consistent with previous reports in ruminants that the immature immune system responds poorly to prior immunisation (33, 34). Further, age differentially affected the ability of sheep to control worm numbers and the numbers of parasite eggs produced in their faeces: while we observed a significant reduction of parasite burden in 6-month-old animals irrespective of their vaccination status, age alone did not significantly affect total egg output (Fig. 1). This suggests that maturation of the immune system allows better control of *T. circumcincta* worm burden with age but may have a more limited effect on parasite transmission via parasite egg shedding. Such apparent compensation and density-dependent fecundity in parasites have been previously observed in this (35) and other host-parasite systems (36–38). Further, while the prototype vaccine reduced parasite and egg densities, it did not totally eliminate the presence of high worm egg shedders, raising the possibility that inherently “ wormy” individuals may not respond well to vaccination. Reducing infection in these poor vaccine responders is of particular importance for limiting the spread of infection in vaccinated flocks.

In this study, we took a novel systems vaccinology approach to identify immune transcriptional pathways that predict both post-mortem worm burdens and cFEC in vaccinated individuals. While similar approaches have been taken to understand the mechanisms by which vaccines stimulate protection, these have largely focused on repeated analysis of blood leukocyte transcriptomes which are clustered into blood transcriptional modules (BTM) and subsequently correlated with antibody or cellular immune phenotypes (39–41). This approach, while providing important information on immune pathways involved in vaccine responses, relies on capturing transcriptomic signatures of recirculating leukocytes which may not truly reflect the immune response at the site(s) of infection or vaccination. In contrast, the application of a gastric cannulation technique in this study allowed repeated sampling of the gastric mucosa, resulting in an unparalleled temporal evaluation of the local transcriptomic responses over time at the parasite’s predilection site. Furthermore, we used a novel machine-learning approach to maximise the generalisability of the qualitative and quantitative associations between immune responses and worm numbers or fecundity. Thus, the combination of repeated *in situ* sampling, unbiased selection of immune mediators of vaccine-driven protection, and pathway analysis, allowed us to generate a biologically-interpretable, robust, and dynamical representation of how vaccination affects the immune response to *T. circumcincta* infection in lambs.

Using this approach, analysis of the immune response at the site of infection revealed that three main events were indicative of interactions between vaccination and parasite life history and were predictive of parasite burdens. First, priming of the immune system by the vaccine prior to challenge: immune pathways that determined the worm burdens measured at day 49 post challenge were already highly activated in the transcriptomes before challenge, likely driven by the pre-challenge vaccination. Second, the response to parasite life history events: while reduced pathway enrichment observed between days 7 and 28 suggests a down-regulation of immune activity in the abomasum once founder populations of parasites had established infection, we observed an activation of more directed responses to ongoing parasitological events involved in innate responses (e.g. eosinophil CCR3, NF-κB activation and NO/ROS expression by macrophages) and antigen-processing and lymphocyte recruitment and activation (CD40, IL-7, CXCR4 signalling) 1–2 weeks post-challenge, matching the timing of the emergence of the first larvae from the gastric glands (42). This was followed by inflammatory and stress responses (e.g. IL-17 signalling, TLR-signalling, glucocorticoid receptor signalling) around 3 weeks after infection when parasites begin producing eggs (see Fig. 2). And third, the tighter correlations between gene expression levels and parasite numbers at the last sampling time-point immediately prior to post-mortem analysis of worm burdens (day 42) point to confounding between the effects of vaccination and the temporal proximity between immune tissue sampling (day 42) and measurement of parasite counts (day 49).

Further investigation therefore focused on three time-points, days 0, 7 and 21, which were most predictive of parasite numbers and/or fecundity, to identify pathways that were associated with vaccine-induced protection. Of most interest were protective pathways enriched in the mucosa of vaccinated lambs immediately prior to challenge. These included Th1, IL-6 and interferon signalling pathways which negatively correlated with worm burdens, and IL-17A, IL-9, CCR3 and TGF-β pathways associated with lower parasite egg output, and suggested that the systemically delivered vaccine was able to modulate the immune system at a distant mucosal site. This was a rather surprising observation, as while immune signatures have previously been reported in peripheral blood, for example following yellow fever and influenza vaccination (40, 43), systemically delivered vaccines are generally considered poor at inducing mucosal immune responses due to the inability of the vaccines to induce appropriate homing receptors on activated lymphocytes (44). Furthermore, while some adjuvants (e.g. TLR agonists and bacterial ADP-ribosylating toxin adjuvants) have been shown to confer mucosal homing properties (45), this has not been widely reported for saponin-based adjuvants such as Quil-A used here, although our results are consistent with an earlier study of a systemically delivered *Ostertagia ostertagi* subunit vaccine formulated with Quil-A that reported similar mucosal priming of immune cells, and in particular natural killer (NK) cells (46).

Regardless of the mechanism by which this mucosal priming operates, the vaccine appears to promote a protective Th1/Th17 type response within the mucosa, with evidence of active Th17 polarisation potentially via IL-6 and TGF-β (47–49), as well as evidence of enrichment of Th2-associated pathways, CCR3 and IL-9, more established effectors of anti-parasite immunity (50), possibly via MIF, which was recently reported as essential to type 2 immunity against *Heligmosomoides polygyrus* in mice (51). Interestingly, some of these pathways (MIF, IL17A, and CCR3) were also associated with reduced parasite egg output in control lambs, suggesting that of the protective pathways induced by the vaccine, anti-worm Th1 responses were most unique. The association of vaccine-induced protection with Th1 immunity was maintained at day 7 post-infection, with Th1 pathways being associated with both worm burdens and egg output. By 21 days, a wider range of protective pathways were identified, including those associated with tissue injury and inflammation. At this time-point, both Th1 and Th2 signalling pathways were associated with protection, indicating a broadening of the immune response.

While these associations will require further causal validation, they suggest that for optimal protection, the vaccine may prime the mucosa towards a Th1 and potentially Th17-type response early in infection before broadening out to a more mixed Th1/Th2 response. This is potentially contentious, as it is well-established in numerous animal models that protective immunity to gastrointestinal parasites, including *T. circumcincta*, is associated with Th2 immunity, whereas Th1 and Th17 immunity is associated with susceptibility (52–56). However, this appears to be a rather simplistic model based on counter-regulation of Th immune responses derived from tightly-controlled laboratory models and/or analysis of responses at limited time-points post-infection. Indeed, our longitudinal sampling at the site of parasite infection coupled with whole-transcriptome analysis is likely to have allowed a finer description of the sequence of protective responses, revealing contributions to the protective response of Th1, Th17 and Th2 responses at different phases of the infection within the same individual. These results are consistent with a previous study in sheep, in which natural resistance to *T. circumcincta* was associated with early Th1 responses prior to development of Th2 immunity (57), and a more recent study in which early activation of the Th17 pathway was associated with natural resistance to *Haemonchus contortus* in goats (58). The role of Th17 in protection is potentially explained by the ability of this pathway to elicit innate lymphoid cells and multipotent progenitor type 2 cells early in infection that subsequently promote CD4^+^ Th2 cells and associated cytokine expression (59–61). This then activates antiparasitic effector cells such as eosinophils (62, 63), consistent with our finding that CCR3 signalling in eosinophils at D21 post challenge was negatively associated with egg output (Fig. 3). The previous association of Th17 responses with susceptibility was determined after more long-standing (12 week) *T. circumcincta* infection (56), where the association could be explained by triggering of Th17 responses secondary to gut barrier disruption (64).

Further investigation into how vaccination and age affected the expression of these immune pathways revealed that 6mo lambs increased the expression of the pathways to a greater level than did 3mo lambs at D21, which coincides with the first emergence of parasites from gastric crypts into the lumen of the abomasum (29, 30). This age-dependant expression of protective immune pathways largely involved the activation and maintenance of the adaptive response. This late maturation of the adaptive response to helminths observed in lambs is consistent with reports of later maturation of T cells in sheep (65, 66) and in other host-parasite systems as previously reported (26). Conversely, innate pathways appeared age-independent for these two cohorts. However, vaccination alone significantly explained the upregulation of innate pathways such as those involved in acute phase response signalling, LPS/IL-1 mediated inhibition of RXR functions, TLR signalling, and MIF regulation pathways, only a subset of which are reported to be enhanced by saponin-based adjuvants (67, 68), the others likely induced by the infection.

In conclusion, most protective-associated pathways were induced by vaccination and in older animals; the immune pathways found to control adult worms and eggs only partially overlapped and were activated at different phases of the challenge infection, indicating the need for anthelmintic vaccines to stimulate a broad set of pathways, rather than just antibody production alone; protective pathways enriched pre-challenge in vaccinates suggest specific adjuvants, such as those promoting Th1/Th17 responses may be useful to improve vaccine performance.

## Methods

### Lambs & Infection

Texel cross lambs were randomly allocated to four groups which were balanced for weight and sex. The groups comprised vaccinated 3-month-old lambs (n = 15), control (adjuvant only) 3-month-old lambs (n = 16), vaccinated 6-month-old lambs (n = 15), and control (adjuvant only) 6-month-old lambs (n = 16). All lambs were infected with 2,000 *T. circumcincta* L3 stage larvae three times per week for four weeks beginning on the day of the final immunisation.

### Prototype vaccine formulation

Each lamb in the vaccinated groups was injected with 400 µg of a recombinant protein mix as described previously (19), containing 50µg of each protein. PBS soluble proteins Tci-ASP-1, Tci-MIF-1, Tci-TGH-2, Tci-APY-1, Tci-SAA-1, Tci-CF-1 and Tci-ES20 were administered as a single injection with 5mg Quil-A (Brenntag Biosector). Tci-MEP-1 was administered separately in PBS with 2M urea and 5mg Quil-A. Injections were given subcutaneously at two sites on the neck. Control animals received injections containing PBS/urea and Quil-A only. Lambs received three immunisations at three-weekly intervals (day 0, 21, 42) with the first immunisation at 3 or 6 months of age (Fig. S1).

### Sampling

Seven animals in each vaccine group and eight animals in the control groups were fitted with abomasal cannulae, as previously described (69), to allow repeated biopsy of the abomasal mucosa throughout the trickle infection period. Briefly, peri-operative analgesia was administered prior to anaesthetic induction using Meloxicam (Metacam®, Boehringer Ingleheim) at 1 mg/kg body weight (BW). Anaesthesia was induced by intra-venous Propofol (PropoFlo™, Zoetis) at 3mg/kg BW and maintained using Isoflurane (IsoFlo™, Zoetis). Following surgical preparation of the site, the abomasum was located and exteriorised. Abomasal cannulae, constructed from a modified disposable 10 ml syringe barrels (PlastiPak™, Becton and Dickinson) (69), were fitted midway between the mesenteric border and the greater curvature of the lateral wall, approximately 7 cm cranial to the pylorus. The free end of the cannulae were exteriorised through a laparotomy incision in the abdominal wall, then anchored using external neoprene flanges. Surgery to fit the cannulae was performed between the second and third immunisation time-points. Over 49 days, abomasal biopsies were taken using a pair of 30 cm long 5 mm × 2 mm punch mucosal biopsy forceps (Richard Wolf GmBH), inserted via the cannulae as follows: three biopsies per animal at each time-point on days 0, 7, 14, 21, 28, 42, and 49 (Fig. S1). The three biopsies per animal were taken in a clockwise manner to ensure different sites of the abomasal mucosa were sampled. Samples were placed immediately into RNAlater (Sigma-Aldrich) and stored at -80 °C for subsequent RNA extraction.

### RNA-Seq library preparation and sequencing

Total RNA was extracted from the abomasal biopsies as follows: the 645 abomasal biopsy samples were homogenised in RLT buffer (Qiagen Ltd, UK) using a Precellys bead basher (Bertin Instruments, UK) with CK28 bead tubes (Stretton Scientific, UK). Samples were centrifuged at 14,000 g at 4°C for 10 mins and the supernatant collected for processing using a RNeasy mini-isolation kit (Qiagen Ltd, UK) according to the manufacturers’ protocol, including an on-column DNase digestion. RNA quality and integrity were assessed using a Nanodrop spectrophotometer (Thermo Fisher, UK) and a Bioanalyser RNA Nanochip (Agilent Technologies Ltd, UK). The yield of total RNA was determined on a Qubit Fluorometer (Thermo Fisher, UK) using the Broad Range RNA kit (Thermo Fisher, UK). The RNA isolated from three biopsies per animal for each time-point was pooled 1:1:1 by weight to generate the final samples for RNA-seq assessment. The resulting 180 RNA samples were sequenced on an Illumina NextSeq 500 by Glasgow Polyomics (70) generating 75 bp paired-end reads at an average sequencing depth of 25 Mbp/sample.

### RNA-Seq quality control and alignment

The sequencing quality controls were finished by FastQC (v0.11.5), which provides a comprehensive report for each RNA-Seq sample. Base calls were made using the Illumina CASAVA 1.8 pipeline. All 180 samples passed the QC filters (MultiQC (71) report, Supplementary Data 1), suggesting that RNA extraction and subsequent sequencing were of good quality and cutadapt (v1.11) was used for adapter trimming. Pseudo alignment of the read data to the latest version of the sheep transcriptome (cDNA) (Oar-v3.1) was performed with Kallisto v0.46.2 (72), generating read count data for each transcript across all samples. The R package tximport (73) was used to prepare the abundance matrix for downstream analysis. All samples were normalised using DESeq2 (74) and genes with read counts greater than five across at least two samples were selected for downstream analysis.

### Gene network analysis and machine learning

We first evaluated whether any structure in gene expression could be visualised using t-Distributed Stochastic Neighbour Embedding (t-SNE). To reduce the dimensionality of the transcriptome, we then used the R package Weighted Correlation Network Analysis (WGCNA) (27) to generate gene co-expression networks from vaccinated groups (3mo-Vax and 6mo-Vax) at six time-points after start of the challenge (Days 0 – 42), using the full transcriptome. WGCNA groups genes and builds networks using the co-expression similarity measure defined as 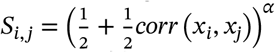, where *S*_*i,j*_ is the correlation between gene expressions *x*_*i*_ and *x*_*j*_, and *α* is the soft threshold weight selected by scale-free topology criterion (27, 75) set at 6 (Day 0), 8 (Day 7), 7 (Day 14), 12 (Day 21), 8 (Day 28), and 9 (Day 42). The eigengene of each cluster was used to quantify its overall expression. To select the genes which best predicted the parasitological read-out of interest (i.e., worm burdens and cFEC), we used the ElasticNet algorithm (76) from Python’s Scikit-Learn software library to fit a linear regression between the eigengenes and worm burdens or cFEC, and then ranked the gene clusters by their resulting coefficients. All WGCNA modules for which the ElasticNet coefficient was not null were retained for further pathway analysis.

### Pathway analysis

Pathway enrichment was generated with Ingenuity Pathway Analysis (IPA, QIAGEN Inc.) (28) in which each gene identifier was mapped to its corresponding gene object in Ingenuity’s Knowledge Base using canonical pathway analysis to identify the biological pathways of most significance. Only immune pathways among those identified in IPA (Supplementary Data 2) were retained for further analysis.

### Time series differential gene expression analysis

The R package maSigPro (77), a two-step regression to find significant differences between treatments over time, was used for gene differential expression analysis with multiple time-points (Supplementary Data 3).

### Statistical methods

The effects of age and immunisation on both worm burdens and cFEC were assessed with generalised linear models for negative binomial distributions with a log link, and residuals tested for normality using the Jarque-Bera test for normality. The effects of age, immunisation, and their interaction on the eigengene of each immune pathway were assessed with linear mixed models using sampling date as a random effect to account for repeated sampling. The t statistic was used to indicate effect sizes as fold deviation between group means (31).

## Acknowledgements

We thank Leigh Andrews, Alison Morrison and Dave Bartley, Moredun Research Institute, for their help in conducting animal studies and in the provision of parasite material and the Bioservices Unit, Moredun Research Institute, for expert care of the animals. We also thank Manus Graham and Roy Davie, Moredun Research Institute, for assistance with gastric cannulation surgeries. This work was supported by the BBSRC (Grant references BB/M012956/1 and BB/M011968/1).

## Supplementary information

**Supplementary Data 1**

`multiqc-report-biopsy.html`: Quality check report of 180 RNA-seq sample FastQC reports using MultiQC (71).

**Supplementary Data 2**

`ipa_3V_6V.ods`: Ingenuity Pathway analysis (IPA) results for all the WGCNA clusters shown in Fig. 2.

**Supplementary Data 3**

`gene-cluster-masigpro.ods`: Differentially expressed gene list between the age groups. The expression levels are shown in Fig. S5C.

## Supplementary Figures

### Study design

**Fig. S1.**
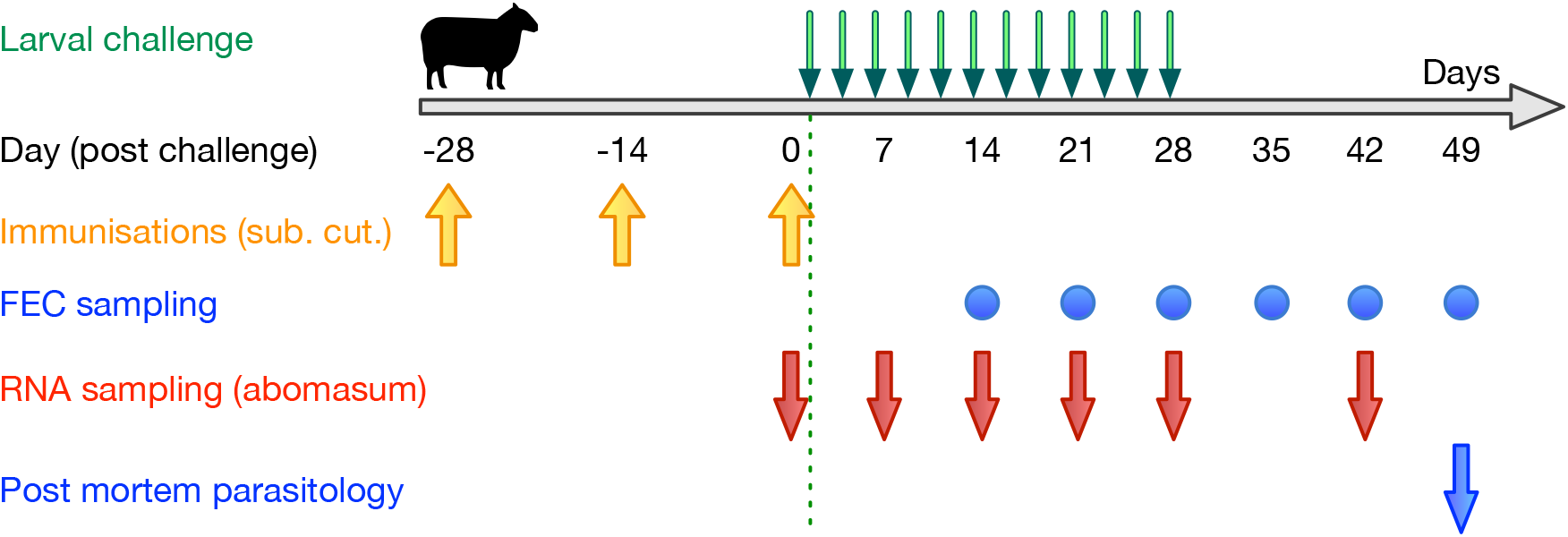
Experimental design. Timeline of immunisation, challenge, and sampling of sheep.

### Methods flow overview

**Fig. S2.**
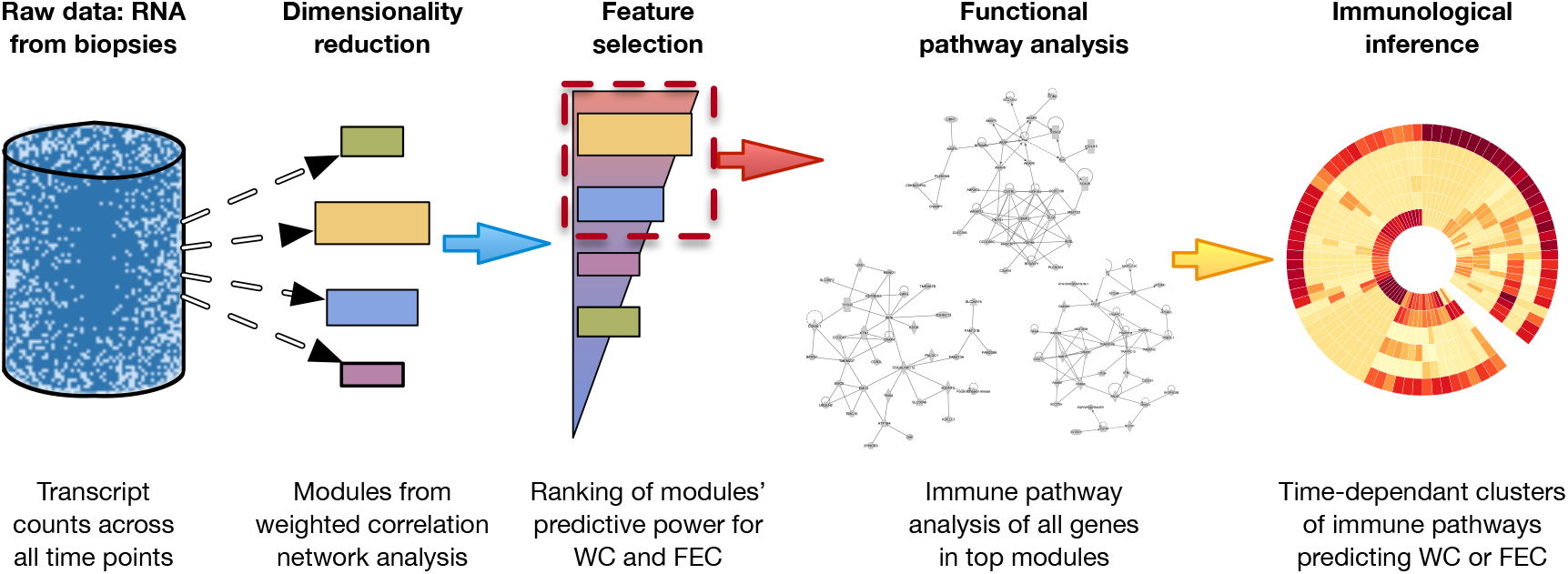
Data analysis pipeline overview. Depiction of the flow through which raw transcript counts were taken to extract immunological information relevant to the response to vaccination and infection.

### Clustering of all transcriptomes using t-SNE

**Fig. S3.**
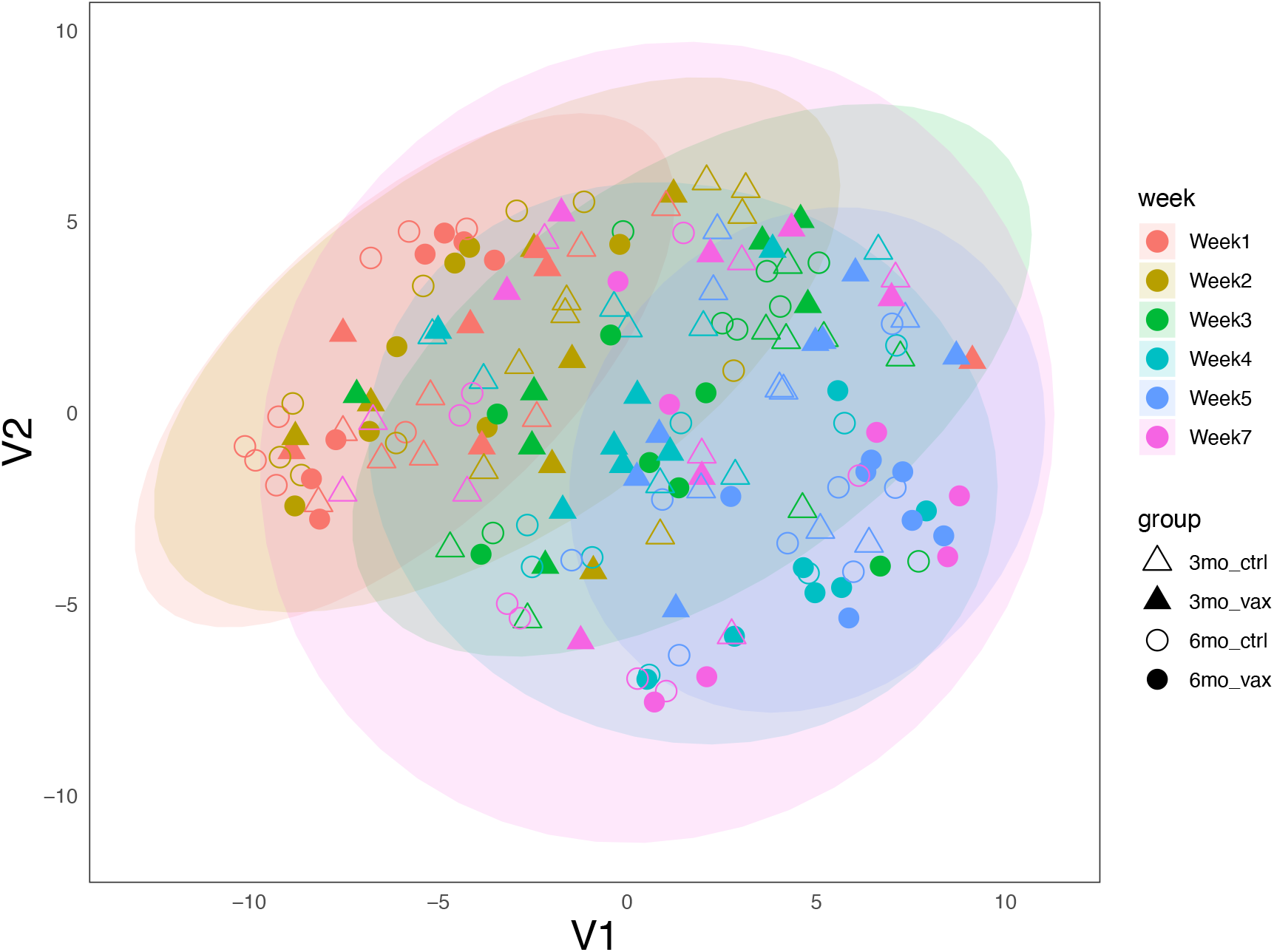
Unsupervised clustering of transcriptomes per treatment and day post challenge. The ∼15K transcripts were reduced to two components using t-Distributed Stochastic Neighbour Embedding (t-SNE) (78) to visualise the datasets in two-dimensional space. Samples clustered weakly by treatment but more distinctly by day post challenge (DPC), specifically before *vs*. after 14 DPC.

### Weight of WGCNA modules predicting parasite burdens

**Fig. S4.**
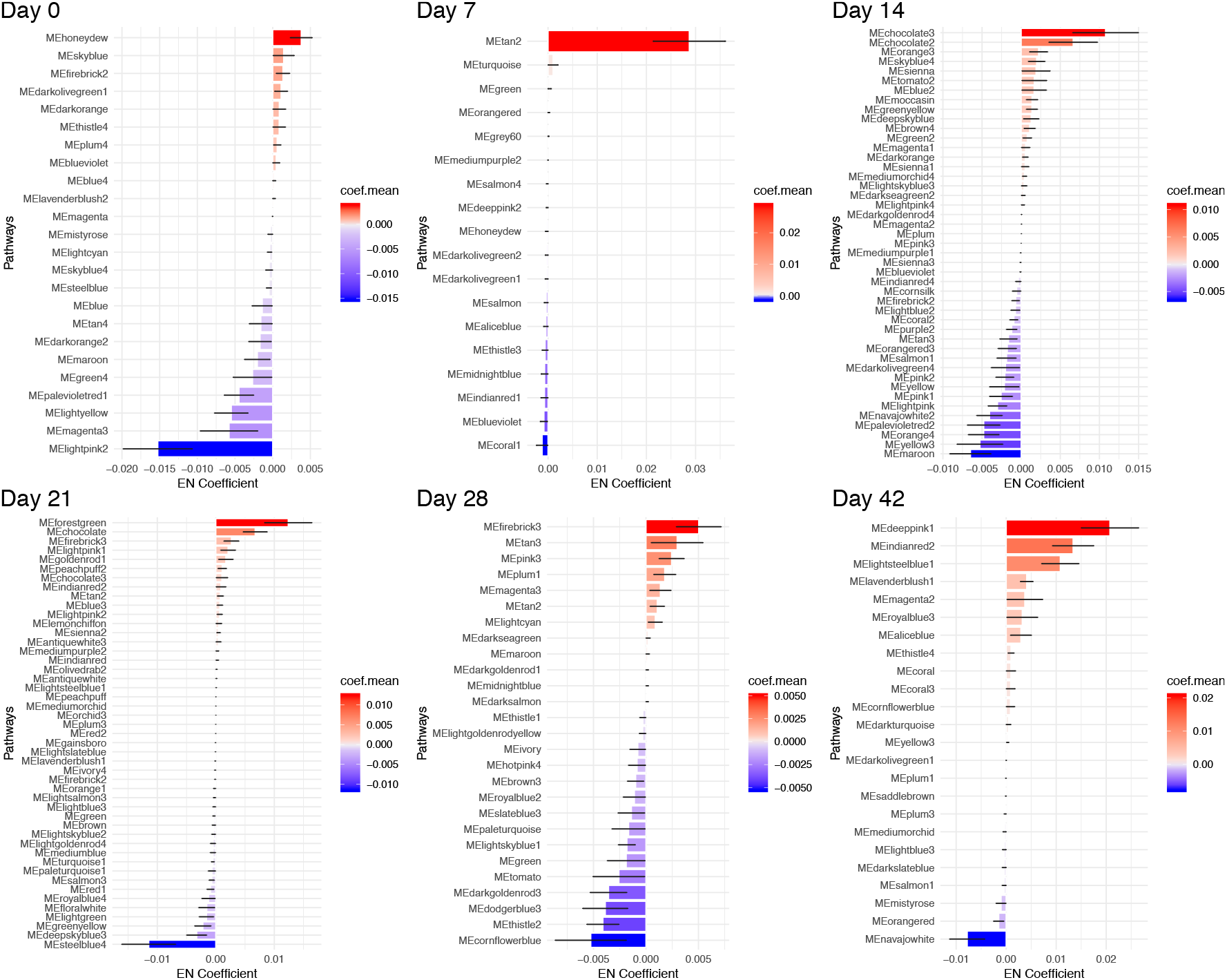
ElasticNet coefficients bar plot of WGCNA modules associated with worm burden and cFEC across 42 days post challenge. Only modules with ElasticNet coefficients ≠ 0 are depicted and were retained for further analysis.

### Differential expression analysis of genes included in WGCNA modules

**Fig. S5.**
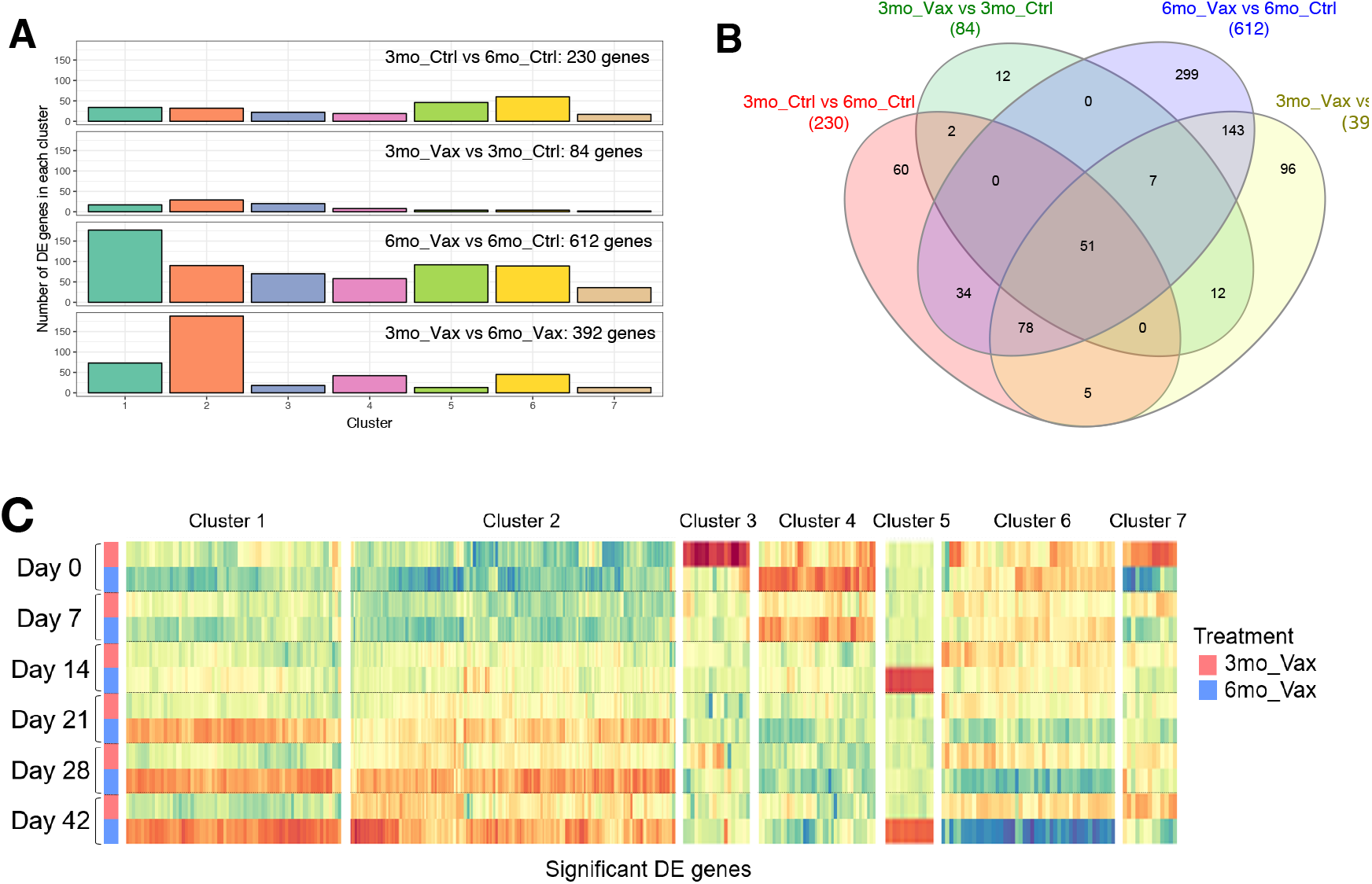
Time dynamics of differentially expressed (DE) genes. (*A*), Number of DE genes in each WGCNA modules within each of 4 pairwise DE pairwise comparisons: 3 month-old control *vs*. 6 month-old control (‘3mo-Ctrl vs 6mo-Ctrl’), 3 month-old vaccinated *vs*. 3 month-old control (‘3mo-Vax *vs* 3mo-Ctrl’), 6 month-old vaccinated *vs*. 6 month-old control (‘6mo-Vax *vs* 6mo-Ctrl’) and 3 month-old vaccinated *vs*. 6 month-old vaccinated (‘3mo-Vax *vs* 6mo-Vax’). (*B*), Venn diagram of all genes from four pairwise comparisons. (*C*) Heat map of the temporal expression patterns of significant DE gene expression. Individual genes within clusters 1–7 are shown in columns (see full gene list in Supplementary Data 2). Each row represents either 3mo or 6mo vaccinated lambs at different time points.

### Time-course of immune pathways significantly represented in abomasal transcriptomes

**Fig. S6.**
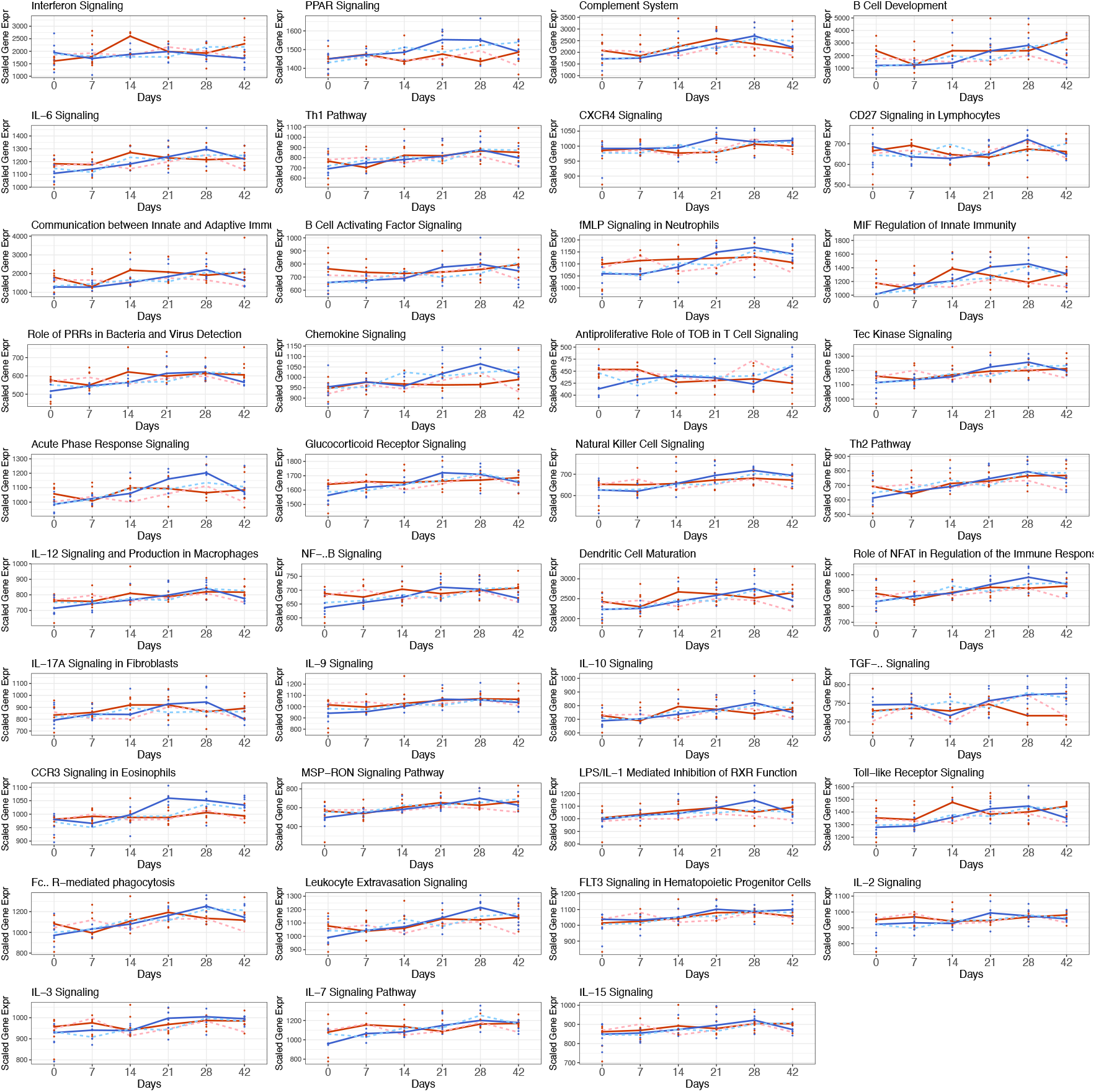
Time course of pathways significantly represented in abomasal transcriptomes. Mean scaled expression levels in each pathway predictive of worm burdens and/or cFEC, selected in Fig. 2. Points represent individual lambs at each time point and lines represent corresponding mean values for: vaccinated (solid lines); control (dashed lines); 3-month-old (red); and 6-month-old (blue) lambs.

